# Multiplex single-molecule DNA barcoding using an oligonucleotide ligation assay

**DOI:** 10.1101/265215

**Authors:** Ivo Severins, Malwina Szczepaniak, Chirlmin Joo

**Affiliations:** Kavli Institute of NanoScience, Department of BioNanoScience, Delft University of Technology, Delft, 2629HZ, The Netherlands

**Author notes:** To whom correspondence should be addressed.; (CJ), (MS). The authors wish it to be known that, in their opinion, the first two authors should be regarded as joint First Authors.

## Abstract

Detection of specific nucleic acid sequences is invaluable in biological studies such as genetic disease diagnostics and genome profiling. Here we developed a highly sensitive and specific detection method that combines an advanced oligonucleotide ligation assay (OLA) with multicolor single-molecule fluorescence. We demonstrated that 7-nt long DNA barcodes have the optimal short length to ascertain specificity while being long enough for sufficient ligation. Using four spectrally separated fluorophores to label DNA barcodes, we simultaneously distinguished four DNA target sequences differing by only a single nucleotide. Our new single-molecule approach will allow for accurate identification of low abundance molecules without the need for target DNA pre-amplification.

## INTRODUCTION

Personalized medicine relies on recognition of specific nucleic acid sequences. Variations in DNA sequences are associated with risks of developing diseases and with varying metabolic response to drugs or vaccines. Many of the currently available nucleic acid recognition methods rely on DNA sample amplification that involves polymerase chain reaction (PCR) (1–4). Even though PCR revolutionized life sciences research, this method is not completely error-free and may lead to a false positive detection of mutations (5). Thus, it has become necessary to develop a method to detect even single-nucleotide variations without the need to employ a step that could be a possible source of an erroneous readout. In addition, such a technique should be sensitive enough to detect low abundance target molecules.

DNA hybridization and specificity of enzymatic ligation (6) of two complementary nucleic acid fragments constitute two basic elements of many bulk and single-molecule nucleic acids detection methods (for more exhaustive reviews of the methods used in single nucleotide genotyping, see (7-9)). For example, Tyagi et al. (10, 11) designed molecular beacons that exploit the hybridization capability of DNA. These hairpin-shaped oligonucleotide probes, labelled with an internally quenched fluorophore, were used to distinguish four different target DNA strands in bulk in homogenous solution. In independent studies, Landegren et al. (12) and Alves et al. (13) combined the annealing property with the enzymatic ligation reaction, thus introducing the oligonucleotide ligation assay (OLA). This original ligation assay was later combined with a Förster resonance energy transfer (FRET) detection method (14). Similar to OLA, padlock probes proposed by Nilsson et al. (15), take advantage of both reactions, hybridization and ligation, creating circular DNA molecules catenated to the target sequence.

More recently, in an attempt to increase sensitivity and specificity without the need for enzymatic target amplification, researchers moved towards single-molecule methods. Castro et al. (16) combined DNA hybridization with laser-based single-molecule detection of single-copy genes in a complex genome. Also, molecular beacons were extensively studied and their various versions were employed in the single-molecule SNP genotyping techniques. Single-molecule FRET allowed to analyze low abundancy point mutations in K-ras oncogenes by developing reverse molecular beacons (17). Another variation of molecular beacons called “Smart Probes” was developed to minimize unwanted background signal thus increasing identification sensitivity (18). Wang et al. (19) introduced molecular confinement via electrokinetic focusing, which, coupled with the original molecular beacons and a confocal fluorescence spectroscope, brought the limit of detection down to attomolar range. Similar detection sensitivity, attributed to even lower background signal, was shown by Zhang et al. (20, 21) for a quantum dot-FRET nanosensing platform. Most of the single-molecule techniques utilize fluorescence signal for detection. A distinct detection method (22) relies on the measurement of the extension of a DNA hairpin attached to a magnetic bead, which is controlled by a magnetic trap.

However, many of the existing single-molecule approaches use low volume detection methods (e.g. confocal microscopy), which limit the resulting data yield. Moreover, they often apply relatively long DNA probes (15 nt or more) that can potentially lead to false positive detection. The only so-far reported technique that uses shorter, and therefore more specific probes employs a magnetic trap, a more specialized and less common detection method than fluorescence.

Here we furthered the bulk OLA technique (12, 13) by combining it with a single-molecule fluorescence detection scheme. Our method provides a highly specific and sensitive DNA barcoding tool with potential for multiplexing applications. The high specificity is assured by the use of a pair of very short 7-nt DNA barcodes (probes), which are ligated only if complete complementarity to two adjacent sites on a target DNA molecule is realized. The sensitivity, on the other hand, is provided by the single molecule approach, which enables the detection of low abundance targets. We show that our method can be applied to simultaneously distinguish at least four single-nucleotide variants.

## MATERIALS AND METHODS

### Slide preparation

Microfluidic chambers used in all DNA barcoding experiments were prepared on the polyethylene glycol (PEG) coated quartz microscopy slides, according to a previously published video protocol (23). As in the protocol, a fraction of the PEG was biotinylated to enable DNA immobilization. The quality of the slide surface was further improved by 10 min incubation with 5% (v/v) Tween-20 (Sigma-Aldrich, St. Louis, MO) in T50 buffer (10 mM Tris-HCl [pH 8.0], 50 mM NaCl), followed by an extensive wash with T50 (24).

### Target DNA immobilization

As target for barcoding, 60-nt long single-stranded DNA molecules were used (Integrated DNA Technologies, Coralville, IA and ELLA Biotech, Martinsried, Germany), which were annealed to an 18-nt biotinylated anchor sequence (sequence information can be found in Table S1 in the Supporting Material). Target strands were immobilized through streptavidin-biotin conjugation of the biotinylated anchor DNA and the biotinylated PEG coating. This was achieved by incubating the slide surface with 0.1 mg/mL streptavidin (Invitrogen, Carlsbad, CA) for 1 minute, followed by a 3-minute incubation with 38 pM target DNA; when multiple targets were used the total target concentration remained 38 pM. Between incubations the channels were flushed with T50.

### Barcode preparation

Short DNA oligonucleotides (Integrated DNA Technologies and ELLA Biotech) were deployed as barcodes complementary to two neighboring sites on the target DNA. The barcodes were termed upstream if they were complementary to the 5’-site and downstream if they were complementary to the 3’-site on the target sequence (see Fig. 1). To enable ligation, all upstream barcodes had a phosphorylated 5’-end, while all barcodes contained one amine modification (an amino group on a six carbon spacer arm) for fluorescent labelling, either at an internal thymine or at the 3’- or 5’-end. Barcodes were labelled with Alexa Fluor 488 (Invitrogen), Cy3, Cy5 or Cy7 (GE Healthcare, Little Chalfont, United Kingdom) via N-Hydroxysuccinimide ester crosslinking. Free dye was removed through ethanol precipitation.

**FIGURE 1.**
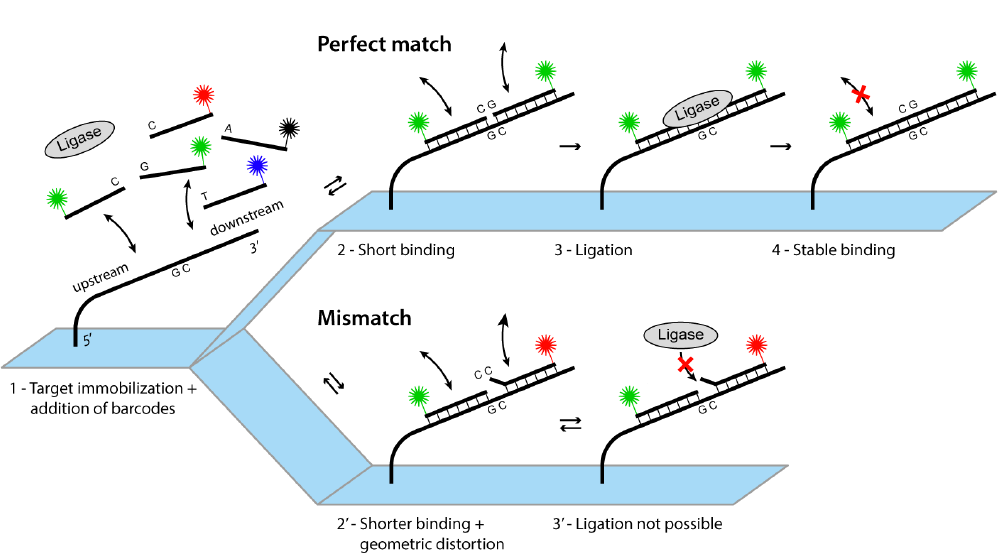
DNA barcoding experimental scheme. Target DNA strands are immobilized on a microscope slide and dye-labeled barcodes are introduced together with T4 DNA ligase in the microfluidic chamber (1). Complementary barcodes bind shortly to the target site (2), while mismatched barcodes bind on an even shorter time scale (2’). Successful ligation is observed for the complementary barcodes (3), but not for the mismatched barcodes (3’). Ligation product shows stable binding to the target DNA (4), while mismatched barcodes dissociate and are washed away before imaging.

### Barcoding procedure

Immobilized target DNA was incubated with 50 nM of each upstream and 50 nM of each downstream barcode (independent of the number of barcodes used) and 14 Weiss units/mL of T4 DNA ligase (Thermo Fisher Scientific) in freshly prepared ligation buffer (40 mM Tris-HCl [pH 7.6], 10 mM MgCl2, 10 mM dithiothreitol, 0.5 mM ATP) for 1 hour at 25 °C. Subsequently, the ligation buffer was replaced by imaging buffer (20 mM Tris-HCl [pH 8.0], 50 mM MgCl2, 50 mM NaCl, 1 mM Trolox, 0.1 mg/mL glucose oxidase, 17 μL/mL catalase, 0.8% w/v glucose) to enhance photostability of the dyes during imaging (25). For four-color imaging (see below) buffer components were dissolved in deuterium oxide instead of water to increase fluorophore brightness (26).

### Restriction reaction

Restriction of target-attached barcodes at the formed GGCC palindromic sequence was performed by incubation with 30 units/mL HaeIII (New England Biolabs, Ipswich, MA) in CutSmart buffer (20 mM Tris acetate [pH 7.9], 50 mM potassium acetate, 10 mM magnesium acetate, 100 μg/mL BSA; New England Biolabs) for 45 minutes at 25 °C.

### Single-molecule fluorescence microscopy

Image acquisition was performed using a prism type total internal reflection fluorescence (TIRF) microscopy setup. Alexa Fluor 488, Cy3, Cy5, and Cy7 fluorophores were excited with 473 nm (blue), 532 nm (green), 637 nm (red), and 730 nm (near-infrared) lasers (OBIS 473 LX 75 mW, Sapphire 532 LP 100 mW, OBIS 637 LX 140 mW, OBIS 730 LX 30 mW; Coherent, Santa Clara, CA), respectively. The laser beams were combined using dichroic mirrors with 523, 544, and 652 nm cut-off wavelengths (ZT514rdc,ZT532rdc, ZT640rdc, respectively; Chroma, Bellows Falls, VT). Emitted fluorescence was collected using a 60x NA 1.2 water-immersion objective (Olympus, Tokyo, Japan) mounted on an inverted microscope (IX73, Olympus). The image was additionally magnified (2.5x) using two achromatic doublet lenses with 100 and 250 mm focal lengths (AC508-100-A-ML and AC508-250-A-ML, respectively; Thorlabs, Newton, NJ).

For two-color experiments, the green and red lasers were used at a 3:2 power ratio. Scattered excitation light was blocked using a 488/532/635 nm triple notch filter (NF01-488/532/635; Semrock, Rochester, NY) and the remaining signal was projected onto two halves of an EMCCD camera (iXon+ DU-897D; Andor Technology, Belfast, UK) using a 635 nm dichroic mirror (635dcxr; Chroma). For four-color experiments, alternating laser excitation (ALEX) was performed using blue, green, red, and infrared lasers at a power ratio of 8:4:2:3; these values are proportional to the extinction coefficients of the corresponding dyes. Scattered light was blocked by a long-pass filter with 50% transmission at 482 nm (BLP01-473R-25; Semrock) and three individual notch filters with rejection peaks at 532 nm, 633 nm, and 730 nm (NF03-532E-25, NF03-633E-25, and the custom made ZET730NF, respectively; Semrock). The signal was projected onto the same camera, however now divided into four parts using 540 nm, 635 nm, and 740 nm dichroic mirrors (540dcxr, 635dcxr, 740dcxr, respectively; Chroma).

Images were acquired with homemade software written in Visual C++. For each experiment, 5-20 independent fields of view were imaged. It should be noted that CCD camera images shown in Fig. 2a-d, Fig. S1, and Fig. S2 represent one half of a full-size field of view, while the CCD camera images in Fig. S3 represent only a quarter of a full-size field of view.

**FIGURE 2.**
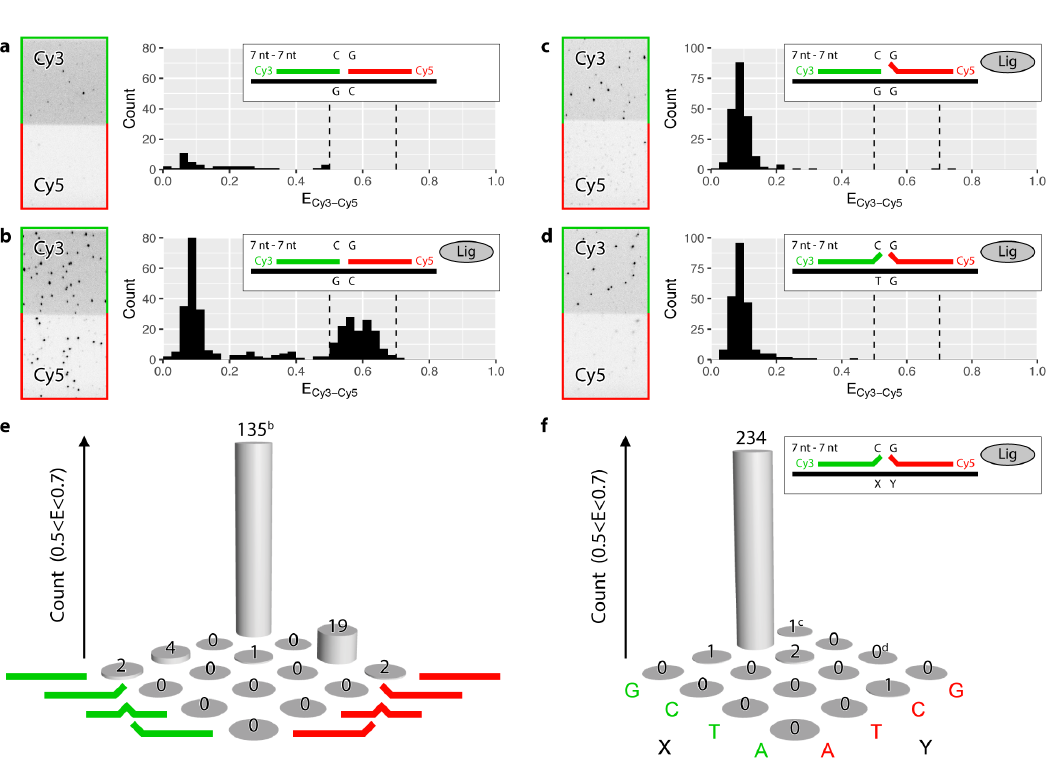
Barcode ligation enables detection of complementary target DNA. (a, b) Detection of DNA target with complementary barcodes in the absence (a) and presence (b) of ligase. (c, d) Detection of DNA target with one (c) or two (d) mismatched barcodes in the presence of ligase. (e-f) Effect of mismatch position within the barcode (at the 5’-end, center, or 3’-end) (e) and base pair identity (f) on the ligation efficiency. Experiments in (a, b, e) and (c, d, f) are based on 5 and 15 fields of view, respectively. Panels (a-d) show CCD camera images (left) of the Cy3 (green) and Cy5 (red) channels upon Cy3 excitation, and experimental schemes and FRET efficiency (E) histograms (right), where the dotted lines indicate the range of E of the target-barcodes complexes (0.5<E<0.7). This range was used to determine the count in the bar plots (e, f); bars indicated with letters “b”, “c”, and “d” were derived from the FRET histograms shown in panels b, c, and d, respectively.

### Data analysis

Single molecules were localized in the acquired images by searching for fluorescence spots with a Gaussian profile and, following background subtraction from the single molecule peaks (27), the intensity time traces were extracted using homemade scripts written in IDL (ITT Visual Information Solutions, Boulder, CO). Time traces were further analyzed using MATLAB (The MathWorks, Natick, MA), where the apparent FRET efficiency (E) and, for four-color data, also stoichiometry (S) were calculated:

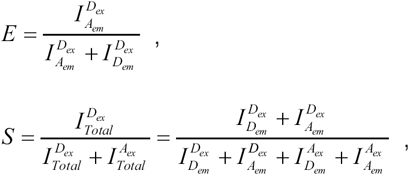

where I denotes the intensity of the donor or acceptor emission (Dem or Aem) upon donor or acceptor excitation (Dex or Aex).

### E and S values in four-color experiments

It is important to take into consideration several factors that can influence the experimentally determined E and S values and cause their deviation from theoretical predictions. First, a non-zero background signal causes a slight shift of E and S toward 0.5; this effect is most prominent for the donor-only and acceptor-only case. Second, the background signal can differ for each laser and each detection channel. Third, E and S values depend on the relative apparent dye intensities, determined e.g. by dye quantum yield and detection efficiency. Fourth, a shift in E and/ or S values may be caused by variations in focus; as refraction indices vary with wavelength, only one of the four dyes can be in focus, leaving the others slightly blurred with reduced intensity. Finally, cross-excitation and spectral bleed-through can contribute to the deviation of E and S from the expected values. In principle, one could computationally correct for many of the above-mentioned factors (28), however this procedure would have to be carefully applied. We found that only adjusting the ratio of different laser powers such that all dyes showed similar apparent intensities upon direct excitation was sufficient for a reliable distinction between the barcode pairs.

### Barcode selection in four-color experiments

To identify the four different barcode pairs among all detected molecules in the four-color experiments, selection criteria based on intensity, stoichiometry and FRET efficiency values were computationally established. As a first step, the intensity histograms were fitted with a sum of univariate Gaussian distributions using the MATLAB function “fit” (see Fig.3,4, S4, and S5), where the number of Gaussians was determined by automatically estimating the number of peaks in the histogram using the MATLAB function “findpeaks”. From these fits, the intensity criteria for each fluorophore were obtained, enabling the separation of the background signal (lowest intensity peak), the single dye peak and the double dye peak. All intensities smaller than the upper boundary of the 99 % confidence interval of the lowest intensity Gaussian were considered background signal. The higher intensity Gaussians corresponded to individual fluorophores, with the selection threshold set to the 95 % confidence interval. Based on intensity, molecules were selected as a specific dye pair (e.g. A488-Cy3) when the fluorescence intensities of the two dyes were larger than the background and within this 95 % confidence interval and, in addition, the remaining fluorophores (e.g. Cy5 and Cy7) had intensities within the background. Subsequently, E-S scatter plots were constructed for the A488-Cy3, Cy3-Cy5, and Cy3-Cy7 pairs. To simplify fitting, in each plot only molecules were included with one or both of the dye pair fluorescence intensities above background and the other two intensities within the background. Next, for each dye-pair the relevant scatter plot was fitted with a sum of three bivariate Gaussian distributions, corresponding to the donor-only, acceptor-only and donor-acceptor populations. This was achieved using the MATLAB function “fit” after first estimating the population locations using the MATLAB function “kmeans”. For selection of the donor-acceptor population a 95% confidence interval for the bivariate Gaussian was used, resulting in an elliptical boundary in the ES-scatter plot (Fig.3, 4, S4, and S5). Selection of the molecules corresponding to the Cy3-Cy3 pair was based on intensity information only. Here, in addition to the background and the single-dye peak, also the higher intensity peak (at approximately twice the single-dye intensity), corresponding to two colocalized Cy3 fluorophores, had to be discerned.

**FIGURE 3.**
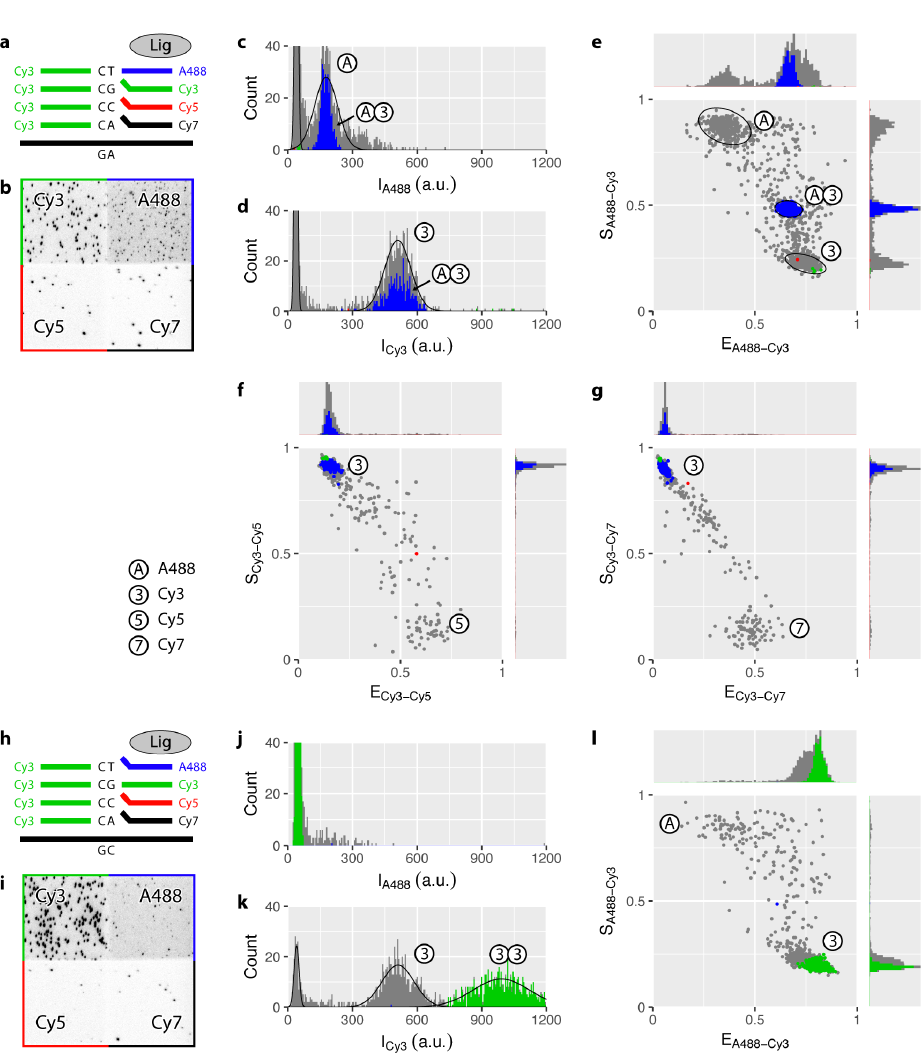
Single target complementary to one of four distinctively labelled barcode pairs. (a, h) Experimental scheme. (b, i) Combined CCD camera image of the four channels: A488 (blue), Cy3 (green), Cy5 (red), and Cy7 (black) upon respective direct excitation. For the original CCD camera images used in (b), see Fig. S8. (c, j) and (d, k) Intensity (I) histogram for A488 and Cy3, respectively. Solid lines show fits of univariate Gaussian distributions. (e, f, g, l) FRET-Stoichiometry (E-S) scatter plots for the A488-Cy3 (e, l), Cy3-Cy5 (f), and Cy3-Cy7 (g) fluorophore pairs. In each plot, only relevant molecules are shown, i.e. with one or both of the two fluorophore intensities above background. Ellipses indicate the 95% confidence interval of fitted bivariate Gaussian distributions. Data based on 20 fields of view. Molecules selected as A488-Cy3, Cy3-Cy3, Cy3-Cy5, and Cy3-Cy7 barcode pairs are indicated with blue, green, red, and black, respectively; unselected molecules are shown in grey. The selection criteria were based on the combination of four experiments, each of them using a different target sequence. Single white circles indicate donor-only or acceptor-only populations, while pairs of white circles indicate donor-acceptor populations, i.e. barcode pairs.

**FIGURE 4.**
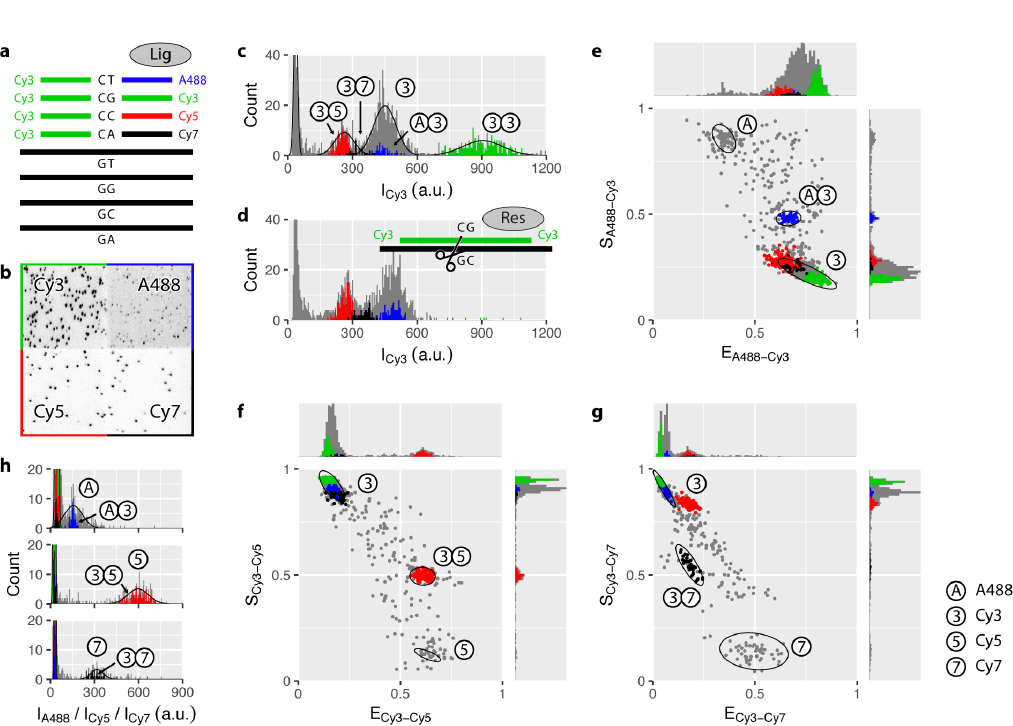
Four targets in mixture identified by four complementary barcode pairs. (a) Experimental scheme. (b) Camera image of the four channels: A488 (blue), Cy3 (green), Cy5 (red), and Cy7 (black) upon respective direct excitation. (c, h) Intensity (I) histograms for Cy3 (c) and A488, Cy5, and Cy7 (h). Solid lines show fits of univariate Gaussian distributions. (d) Intensity histogram for Cy3 fluorophore after addition of a restriction enzyme specific to the bound Cy3-Cy3 barcode pair. (e, f, g) FRET-Stoichiometry (E-S) scatter plot for the A488-Cy3 (e), Cy3-Cy5 (f), and Cy3-Cy7 (g) fluorophore pair. In each plot, only relevant molecules are shown, i.e. with one or both of the two dye intensities above background. Ellipses indicate the 95% confidence interval of fitted bivariate Gaussian distributions. Data based on 20 fields of view. Molecules selected as A488-Cy3, Cy3-Cy3, Cy3-Cy5, and Cy3-Cy7 barcode pairs are indicated with blue, green, red, and black, respectively; unselected molecules are shown in grey. The selection criteria were based on a single experiment with four different target sequences. Single white circles indicate donor-only or acceptor-only populations, while pairs of white circles indicate donor-acceptor populations, i.e. barcode pairs.

In the four single-target experiments, each single experiment provided the selection criteria for only one of the four barcode pairs; therefore, the selection criteria of the four experiments were combined and the combination was subsequently applied to each experiment. Background and Cy3 intensity criteria could be extracted from more than one experiment; in this case conservatively the outermost bounds were used for selection, i.e. the largest background value and the lowest and highest Cy3 intensity (independently). In the four-target experiment, all selection criteria were determined simultaneously.

## RESULTS AND DISCUSSION

### Single-molecule oligonucleotide ligation assay

In a conventional OLA experiment two DNA probes hybridize to two immediately adjacent sequences on a denatured target DNA (12, 13). DNA ligase joins the 3’-end of one probe with 5’-end of the other one only if nucleotides at the junction formed by the juxtaposed strands are correctly base-paired with the target DNA. The newly formed oligonucleotide is de-hybridized from the target DNA and visualized on a separation gel either via autoradiography or fluorescence, provided one of the DNA probes was labelled with 32P or a fluorescent dye, respectively. This method, however, under certain conditions could lead to false positive readout errors, as the length of the used probes (usually 15-20 nt) appears to be sufficient to lead to a stable probe binding to a target DNA, even in the presence of a mismatch. Authors of the OLA technique reported such errors to depend on the ligase and, more importantly, salt concentration (12).

We set out to find an optimal length of the DNA probes, here called DNA barcodes, for our single-molecule fluorescence and FRET experiments. The ideal candidates would be long enough to simultaneously hybridize to the desired site on the target DNA while being sufficiently short to avoid unwanted hybridization to a mismatched target or to a non-targeted part of the sequence. Thus, we designed an assay in which two short DNA barcodes, each labelled with a fluorescent dye, are added with ligase to a microfluidic chamber with immobilized target DNA molecules (Fig.1, steps 1-4). The barcodes are complementary to two adjacent sites on the target strand and should become ligated in the presence of the enzyme. After 1 h incubation time, the extent of ligation can be verified by monitoring FRET efficiency E between the two fluorophores upon excitation of the donor dye. E is defined here as the ratio of the acceptor fluorescence intensity and the sum of the donor and acceptor fluorescence intensities. If both DNA barcodes are simultaneously hybridized to the target DNA, their fluorophores should exhibit FRET, where the FRET efficiency depends on the labeling position (see Table S1 in the Supporting Material for detailed sequences and labeling positions).

We tested 15-nt DNA barcodes, since their length falls in the size range of the original OLA constructs. The experimental scheme shown in Fig. 1 was used here in a two-color fashion, i.e. with only a single upstream and a single downstream barcode labelled with Cy3 and Cy5, respectively. The constructs showed stable binding even in the absence of ligase (as inferred from the observed signal in Cy3 and Cy5 channels in Fig. S1a, as well as the presence of a FRET peak), proving undesirable for further studies. As our aim was to find a set of two DNA barcodes that only after ligation would form a stable duplex with the target DNA, we next evaluated various DNA probes shorter than 15 nt. Our single-molecule data shows that, under our experimental conditions, 7-nt barcodes only transiently hybridize to target sequence in the absence of ligase (Fig.2a) on a time scale shorter than the observation time. Despite such short, limiting interactions, in the presence of ligase we were able to capture stable binding events between a pair of 7-nt barcodes and the target DNA (Fig. 2b). Conversely, replacing one of these DNA barcodes by a 6-nt probe led to a radical decrease in the ligation efficiency (cf. Fig. S1b and S1c). This result is in accordance with the phenomenological rule of seven, postulated by Cisse et al. (29), which states that the annealing efficiency of two complementary single-stranded DNAs into a duplex drastically decreases when the number of contiguous nucleotides forming a duplex changes from 7 to 6. On the other hand, increasing the length of both barcodes to 8 nt resulted in an opposite effect annealing efficiency and thereby ligation efficiency increased in comparison with the 7-nt barcodes (Fig. S2 a-c). However, this increase also facilitated ligation of 8-nt DNA barcodes with introduced single-point mismatches in their sequences, especially when the mismatch was located away from the ligation site. (Fig. S2d-e). Therefore, based on these unwanted false positive read-outs of mismatched 8-nt barcodes and too low detection of 6-nt barcodes, a pair of 7-nt DNA barcodes was deemed a suitable choice for further studies.

### Single-molecule OLA shows single-nucleotide specificity

To further assure the specificity of our assay, single-point mutations were introduced in both 7-nt DNA barcodes. DNA barcodes where mutated at either end (3’ or 5’) or in the center, at the 3rd or 4th nt from the 5’-end (for detailed sequences and labeling positions see Table S1). This resulted in a total of 4 probes per each, upstream or downstream, DNA barcode: one complementary barcode and three mutated ones. We performed 16 independent experiments, in which an equimolar mixture of one upstream and one downstream DNA barcode was incubated in the presence of ligase in a microfluidic chamber with immobilized target DNA (Fig. 1). Counting the number of target DNA molecules with ligated products that showed FRET efficiency between 0.5 and 0.7 (E range for both DNA barcodes complementary to the target strand, Fig. 2b) shows only few successful ligation events that involved one or two mismatched probes (Fig. 2e). Our single-molecule data supports our expectations (Fig.1, steps 1-3’), based on earlier reports on the effect of the mutations at the exact ligation site on the efficiency of the enzymatic reaction (6, 12).

However, we observed one exception: the combination of a complementary downstream DNA barcode with an upstream barcode with the “center” mutation, which presents 19 ligated products as compared to 135 for both barcodes being complementary to the target DNA. This mutation in the center of the upstream barcode (A was replaced by G) most likely led to the formation of a G-T wobble pair, which was postulated half a century ago by Crick (30) and was later shown in a crystal structure of A-DNA, B-DNA, and Z-DNA (31). Brown et al. also concluded that the G-T wobble can be easily accommodated in a DNA double helix without substantial perturbation of its overall conformation. Additionally, the G-T non-canonical pair was found to be the least destabilizing in the NMR in vitro studies (32), while DNA mismatch-repair in vivo studies (33) reported on a G-T mismatch showing the highest repair efficiency amongst all tested mismatches. Thus, we conclude the higher number of ligated products to be a side effect of the specific mutation choice and expect only a negligent number of ligation events, provided the introduced mutation would be of different identity, e.g. if the A was replaced by a T.

Mutations introduced at either side of the ligation junction show the most disruptive effect on ligation efficiency (Fig. 2c); the same effect can be seen when using 8-nt barcodes (see Fig. S2c-e). Therefore, in our experimental design, we decided to probe the target DNA sequence at the ligation site: opposite to the 5’-end of the upstream barcode and 3’-end of the downstream barcode. To ascertain that none of the different mismatch pairs between target DNA and barcode, including potentially formed wobble pairs, give rise to false positive read outs, we measured the extent of ligation for all 16 possible sequences of the target DNA, while keeping the set of DNA barcodes constant throughout all 16 experiments. Once again, only two complementary barcodes became ligated and showed FRET, while introduction of even a single mismatch precluded efficient ligation (Fig.2c, d and f). Notably, a potential wobble G-T pair also did not undergo ligation, despite its supposed structural resemblance to a G-C canonical base-pair. We attribute it to the position at which the wobble pair is present – at the ligation site – where it most likely causes greater steric hindrance as opposed to a wobble pair further away from the ligation site (34, 35). We thus showed that our assay is highly specific to even a single nucleotide mismatch at the ligation site.

To illustrate that our approach of ligating a pair of two 7-nt barcodes has a significant advantage over a simpler assay using a single 14-nt ssDNA probe, we added a single Cy3-labeled 14-nt ssDNA probe, either complementary to the target DNA or containing one or two mutations in its central part, to the microfluidic chamber with immobilized target DNA in the absence of ligase. Comparison of the number of bound molecules shows that a presence of a single mismatch in the middle of the 14-nt probe does not abolish its stable binding to the target sequence (Fig. S3a, b). Moreover, even introduction of two mutations in the 14-nt probe does not completely prevent it from stably binding to the target DNA (Fig. S3c). Therefore, a single 14-nt probe is not recommended as an alternative for ligating two 7-nt barcodes to perform accurate target recognition.

### Four-color detection of four distinct barcode pairs

For the versatile use of our single-molecule method, we increased the number of spectrally distinct fluorophores from two to four by introducing Alexa Fluor 488 (henceforth called A488) and Cy7 to the previously used Cy3 and Cy5. These four fluorophores were attached to one of the four sequence variants of the downstream DNA barcode at its 5’-end, while a single upstream barcode was labeled with Cy3 at its 3’-end. Alternating laser excitation (ALEX) allowed us to cycle through four different laser excitation beams and, for each laser color, to simultaneously collect fluorescence signal in four spectrally separated channels on the CCD detector (Fig. S8).

We set out to characterize each dye pair separately. For this we added a mixture of four downstream DNA barcodes, one upstream DNA barcode, and ligase to a microfluidic chamber with only one of the four complementary target DNA strands immobilized. Based on our two-color experiments (Fig. 2), we expected to observe only the ligation products formed by complementary DNA barcodes. For example, as in one chamber we immobilized target DNA with the GA sequence at the ligation site, we expected to detect mostly a downstream DNA barcode with T at the 3’-end, labeled with A488, and the upstream DNA barcode with C at its 5’-end, labeled with Cy3 (Fig. 3a). Imaging indeed shows that most of the bound DNA barcodes are labeled with either A488 or Cy3, or both (Fig. 3b–d and Fig. S4a).

To identify target DNA molecules with two simultaneously bound barcodes, we used the detected fluorescence intensities to calculate stoichiometry and FRET efficiency that were further combined into scatter plots (36) for each of the three possibly formed FRET pairs (Fig. 3e-g and Fig. S4c). In these scatter plots FRET efficiency E is defined as a ratio of acceptor fluorescence intensity upon donor excitation and the sum of donor and acceptor fluorescence intensities upon donor excitation, while stoichiometry S is defined as the ratio of the sum of donor and acceptor fluorescence intensities upon donor excitation and the sum of donor and acceptor fluorescence intensities upon donor and acceptor excitation. Stoichiometry gives an indication of the presence of donor and acceptor dyes. In an ideal case without e.g. bleed through or cross-excitation between the channels, the donor-only population will theoretically have E = 0 and S = 1, the acceptor-only population will have E = (assuming identical background in both channels) and S = 0, while the donor-acceptor population will have 0 < E < 1 (E value depends on the inter-dye distance) and 0 < S < 1 (with S = 0.5 when equal dye brightness is assumed). The experimental scatter plots (Fig.3e-g and Fig. S4c) indeed show donor-only, acceptor-only and donor-acceptor populations near the expected E and S values. Furthermore, the donor-acceptor populations are found only in the E-S plots of the expected dye-pair given the sequence of immobilized target DNA (e.g. for the GA target sequence donor acceptor populations can be seen only in the A488 Cy3 E-S plot, but not in the Cy3-Cy5 or Cy3-Cy7 E-S plots, see Fig. 3e-g).

For the twin pair Cy3-Cy3, formed in the presence of the GC target sequence (Fig.3h-l), FRET efficiency and stoichiometry cannot be determined; therefore, identification of this pair had to be achieved using fluorescence intensity data alone. The Cy3 intensity histogram (Fig. 3k) shows three peaks; the lowest intensity peak corresponds to the background signal, whereas the remaining two represent one Cy3 labeled barcode and two Cy3-labeled barcodes, respectively, as the latter peak has twice the intensity of the single Cy3 barcode peak. In addition, the intensity histograms of A488 (Fig. 3j), Cy5, and Cy7 (Fig. S4b) show no other signal than the background, further substantiating that barcodes do not bind to a mismatched target sequence. Furthermore, Fig. S4c shows that the Cy3-Cy3 pair does not erroneously contribute to the donor-acceptor populations in the E-S plots of the remaining FRET dye pairs, and therefore it can be reliably discriminated from these pairs.

We next quantified the bound DNA barcode pairs in each of the experiments with one of the four target molecules immobilized and four barcode pairs present in the microfluidic chamber. Selection criteria for each barcode pair were based on Gaussian fits of the intensity, FRET efficiency, and stoichiometry data (for details see Materials and Methods). This procedure yielded 344 A488-Cy3, 681 Cy3-Cy3, 385 Cy3-Cy5, and 259 Cy3-Cy7 barcode pairs bound to their matching targets and only a few percent of the barcode pairs bound to mismatching targets (Table S2), thus confirming the specificity of our method.

### Four-color scheme to distinguish four different DNA target sequences

As a proof of principle, we demonstrated detection of four different DNA sequences in mixture. We introduced the four barcode pairs into a microfluidic chamber with a mixture of the four different immobilized target sequences (Fig. 4a). The camera image (Fig. 4b and Fig. S5a) shows bound molecules in all four detection channels. Intensity histograms and E-S plots (Fig. 4c, e-g and S5b, c) show populations well resembling those observed in the single-target experiments (cf. Fig. 3 and S4). The Cy3 intensity histogram (Fig. 4c), in addition to the peaks representing single-Cy3 and double-Cy3 molecules, shows peaks of Cy3 dyes forming FRET pairs with A488, Cy5, and Cy7 (colored in blue, red, and black, respectively). To quantify the number of bound barcode pairs, we again applied selection criteria based on Gaussian fits (described in detail in the Methods). Using these criteria, 47 A488 Cy3, 226 Cy3-Cy3, 106 Cy3-Cy5, and 35 Cy3-Cy7 molecules were identified.

To verify that the barcodes were correctly bound to their matching target molecules, a restriction enzyme (HaeIII) was applied, which recognizes and specifically cuts the double-stranded GGCC sequence. In our experimental design, this sequence corresponds to the Cy3-Cy3 barcode pair bound to the GC target sequence. In the resulting Cy3 intensity histogram (Fig. 4d), as well as the remaining intensity histograms and E-S plots (Fig. S7), the previously observed Cy3-Cy3 population disappeared, while the other populations remained unchanged. Quantitative analysis, upon applying the same selection criteria as used before adding the restriction enzyme, also shows a dramatic decrease of Cy3-Cy3 pairs to 6, without a substantial effect on the A488-Cy3, Cy3-Cy5 and Cy3-Cy7 pairs (Fig. 5). Additionally, when the same restriction enzyme was used in the single target experiments, again only the Cy3-Cy3 population bound to the GC target sequence vanished, while other populations remained unaffected (Fig. 5 and Fig. S6).

**FIGURE 5.**
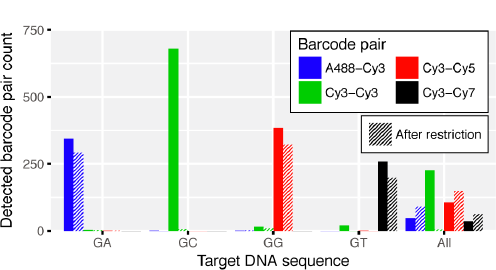
Enzymatic restriction confirms specificity of the DNA barcoding. Number of barcode pairs detected in four-color single-target and four-target experiments, indicat¬ed with the sequence at the ligation site (“GA”, “GC”, “GG”, and “GT”) and with “All”, respectively. Hatched bars show barcode pair counts after addition of restriction enzyme spe¬cific to the bound Cy3-Cy3 barcode pair.

## CONCLUSION

Here we showed that DNA barcoding by ligation of two fluorescently labelled 7-nt single-stranded DNA barcodes, complementary to two neighboring sites on the target DNA strand, can be used to simultaneously distinguish at least four different DNA sequences differing by a single nucleotide. The method relies on a judicious selection of barcode length, which is large enough to allow for simultaneous hybridization of two barcodes leading to an efficient ligation, but is sufficiently small to avoid hybridization and ligation of mismatching barcodes. Barcodes shorter than 7 nt are not readily ligated due to their low binding affinity to target DNA, while barcodes longer than 7 nt allow for their ligation even in the presence of mismatches, thus increasing the risk of detecting false positives.

The inability of the mismatched barcodes to be ligated is the result of a greatly reduced binding affinity in combination with a geometrical distortion in the double helix. This gives the barcoding method its high specificity. The specific location of the mismatch within the barcode and the base pair identity do not appear to be of importance, except when a G-T wobble pair is formed away from the ligation site. Barcodes with a mismatch directly adjacent to the ligation site are least likely to be ligated, due to the additional spatial misalignment of the chemical groups otherwise involved in the ligation reaction.

Distinction of barcode pairs bound to different target sequences can be accomplished by fluorescently labelling each barcode, followed by detection of the different fluorophore combinations. Such detection can be achieved by employing an ALEX excitation scheme coupled with the separate collection of the fluorescence signal from each dye. With the resulting intensity data and the calculated FRET efficiency and stoichiometry, populations of molecules corresponding to the four dye pairs can be visualized. Fitting of these populations with univariate and bivariate Gaussian distributions results in reliable selection criteria that enable computational identification of each dye pair and thereby recognition of four distinct target DNA sequences.

Our DNA barcoding technique exploits inherent sensitivity of a single-molecule approach and therefore may be used as a new method for SNP detection of low abundance target molecules without the need for pre-amplification. Furthermore, its multiplexing feature can be used for example to detect multiple SNPs at once or to simultaneously perform multiple single-molecule experiments. To further increase the multiplexing potential the number of distinguishable barcode pairs should increase. For this, one could take advantage of the remaining fluorophore combinations (e.g. A488-A488, A488-Cy5, etc.). This, together with expanding the set of the upstream barcodes would enable the distinction of a total of ten different barcode pairs and therefore ten different target DNA sequences. Multiple labeling positions leading to various inter-dye distances and thus to different FRET values could be potentially explored. Even though such an expansion of barcode pairs would require an increase in the number of acquired images, it could potentially lead to a much higher number of simultaneously detected target DNA sequences.

## AUTHOR CONTRIBUTIONS

Single-molecule experiments were designed by M.S. and C.J. and performed by M.S. and I.S. Data analysis was done by I.S. I.S., M.S., and C.J. wrote the manuscript. All authors read and approved the manuscript.

## ACKNOWLEDGMENTS

The authors thank members of C.J.’s group for helpful discussions.

